# Systematic comparison of Amber force fields for the mechanical properties of double-stranded DNA

**DOI:** 10.1101/2023.09.25.559335

**Authors:** Carlos Roldán-Piñero, Juan Luengo-Márquez, Salvatore Assenza, Rubén Pérez

**Affiliations:** Departamento de Física Teórica de la Materia Condensada, Universidad Autónoma de Madrid, E-28049 Madrid, Spain; Instituto Nicolás Cabrera, Universidad Autónoma de Madrid, E-28049 Madrid, Spain; Condensed Matter Physics Center (IFIMAC), Universidad Autónoma de Madrid, E-28049 Madrid, Spain

## Abstract

The response of double-stranded DNA to external mechanical stress plays a central role in its interactions with the protein machinery in the cell. Modern atomistic force fields have been shown to provide highly-accurate predictions for the fine structural features of the duplex. In contrast, and despite their pivotal function, less attention has been devoted to the accuracy in the prediction of the elastic parameters. Several reports have addressed the flexibility of double-stranded DNA via all-atom molecular dynamics, yet the collected information is insufficient to have a clear understanding of the relative performance of the various force fields. In this work, we fill this gap by performing a systematic study in which several systems, characterized by different sequence contexts, are simulated with the most up-to-date force fields, bcs1 and OL15, in the presence of external forces with increasing magnitude. Analysis of our results, together with their comparison with previous work focused on bsc0, allows us to unveil the differences in the predicted rigidity between the newest force fields, and suggests a road map to test their performance against experiments.

## 1 Introduction

The elasticity of double-stranded DNA (dsDNA) is a key molecular determinant in the many cellular contexts where this molecule is found, as it affects the binding affinity of dsDNA with proteins^1,2^ as well as the dsDNA response to the mechanical action exerted by proteins^3–7^. This has prompted an intensive research effort in the experimental characterization of the mechanical properties of dsDNA^8–37^. At length scales larger than 10 nm, the mechanics of the duplex is dominated by the persistence length *l_p_*, which regulates the thermally-induced bending of the molecule. At standard ionic strength, *l_p_* lies in the range^8–15^ 45 – 55 nm. At shorter length scales, overall bending fluctuations can be usually neglected and the flexibility of dsDNA is dictated by the stretch modulus *S* and the twist modulus *C*, which account for the deformability of the molecule in elongation and torsion, respectively. These elastic constants are also of interest for large molecules under the presence of mechanical stress, e.g. forces in the range 1 – 50 pN and torques around 1 – 30 pN nm. Quantitatively, it has been found that *S* takes typical values within *S* = 900 – 1600 pN^13–16^, while *C* = 390 – 460 pN nm^2^ ^16–19^. Twist and torsion are also coupled to each other, with dsDNA counterintuitively overwinding upon pulling, as quantified by a negative twist-stretch coupling constant^16,19–21^ −120 pN nm*< g <*−90 pN nm.

All-atom Molecular Dynamic Simulations (AMDSs) are an excellent tool to investigate the mechanical properties of dsDNA, as they enable the inspection of the microscopic mechanisms underlying the global deformation response. For comparison with experiments, the elongation *L* and the torsion *θ* of the molecule as a whole are defined starting from atomistic data, and the elastic constants *S*, *C*, *g* are determined from stress-vs-strain curves^38,39^ or by analyzing the thermal fluctuations of *L* and *θ* ^40,41^. The parm99 force field^42^ in combination with the bsc0 modification^43^ has been shown to predict elastic constants in good quantitative agreement with the experimental estimations^38,39,44–46^. More recently, further force-field modifications have been proposed – OL15^47^ and bsc1^48^ – mainly to improve on the prediction of the helical twist, which was slightly underestimated in bsc0^43^. A thorough comparison of the predicted structural properties has indicated a similar performance of bsc1 and OL15 in capturing the conformational space of dsDNA^49^. Due to their excellent structural agreement with experiments and stability over long simulation times, these modifications have now been accepted as the new standards for simulation of dsDNA.

The values of *S*, *C* and *g* predicted by OL15 and bsc1 have also been found in reasonable quantitative agreement with experiments^40,48,50,51^. Nevertheless, and despite the extensive use of these force fields in the literature, there is a lack of comparative studies highlighting the similarities and differences in the predicted elastic properties within the same sequence context. To our knowledge, only Ref. ^40^ computes the values of *S, C* and *g* by employing bsc1 and OL15 for the same dsDNA fragment, while Ref. ^50^ performs a comparison of the values of *C* obtained for a 32mer via bsc1 and OL15. Future flexibility studies would strongly benefit from a benchmark, where the predictions obtained by the various force fields are assessed in similar conditions and in various sequence contexts, as done previously for the structural features^49^. To meet this need, here we present the results of simulations of various dsDNA sequences performed by using either the bsc1 or the OL15 modifications under the presence of constant pulling forces, and compare them with previous reports employing bsc0 on the same set of duplexes^38,39^. The sequences are chosen so as to encompass all the ten distinct base-pair steps^38,39,45^. We find that, although the three force fields give similar results for *C* and *g*, they can be ranked as bsc0 < bsc1 < OL15 according to the predicted value of *S̃*, with bsc0 providing the most flexible values and OL15 corresponding to the stiffest molecules. Even though in all cases the elastic constants were found to be in reasonable agreement with experiments for a random-like sequence, in line with previous literature^40,48,50,51^, the range of values obtained in the full simulation set suggests that bsc1 might be a preferred option for future flexibility studies.

## 2 Methods

### 2.1 Molecular Dynamics

Following previous work focused on the bsc0 force field^38,39^, we performed AMDSs of the following dsDNA fragments (we write in parentheses the labels that we use in the text to refer to them):

5’-CGCG(AA)_5_CGCG-3’ (poly-AA),
5’-CGCG(AC)_5_CGCG-3’ (poly-AC),
5’-CGCG(AG)_5_CGCG-3’ (poly-AG),
5’-CGCG(AT)_5_CGCG-3’ (poly-AT),
5’-CGCG(CG)_5_CGCG-3’ (poly-CG),
5’-CGCG(GG)_5_CGCG-3’ (poly-GG),
5’-GCGCAATGGAGTACGC-3’ (RNG).

The sequences named poly-XY are obtained as pentamers of the step XY, while the sequence RNG contains all possible base-pair step combinations, and has been introduced in the literature to mimic the behavior of long random sequences employed in the experiments^45^. In all cases, the fragment of interest is sandwiched between two handles in order to minimize end effects^38^.

For each sequence, the dsDNA molecule was built by employing the NAB software within Ambertools19^52^. The system was then placed in a box of side approximately equal to 9 nm and hydrated with explicit water molecules. In order to ensure overall electrical neutrality, a suitable amount of sodium counterions was added to counterbalance the overall negative charge originating from the phosphate-group moieties. For dsDNA, we employed the parm99 force field^42^ with either the bsc1^48^ or the OL15 modifications^47^. Water was modelled according to the TIP3P model^53^, while the Joung-Cheatham parameters were employed for the sodium counterions^54^. Long-range electrostatic effects were accounted for by employing Particle Mesh Ewald, while van der Waals interactions were truncated at the real space cutoff (9 Å). We constrained hydrogen-containing bonds by means of the SHAKE algorithm. The integration timestep was set to 2 fs.

Following standard protocols, the system was first energy-minimized in 5000 steps with restraints applied on the duplex followed by other 5000 steps of unrestrained minimization. Then, the system was heated by linearly increasing the temperature from 0 to 300 K in 300 ps at constant volume. This was followed by an equilibration phase in NPT conditions (*T* = 300 K, *p* = 1 atm) of 20 ns. The last snapshot of the equilibration phase was employed as a starting state for the production simulations, in which the duplex was stretched by a constant force in the NVT ensemble (*T* = 300 K) according to the protocol introduced in Ref.^38^. The force was applied between the geometric centers of C1’ atoms in the sugars within the second and second-to-last base pairs (yellow beads in Fig. 1). For each sequence, five different production simulations were performed at pulling forces equal to 1, 5, 10, 15 and 20 pN. In the production phase, the simulation time was at least equal to 1 µs, leading to a cumulated production time of more than 100 µs. The state of the system was saved every 1000 steps. All simulations were performed with the GPU-accelerated program pmemd.cuda in the AMBER18 suite^52^.

**Figure 1:**
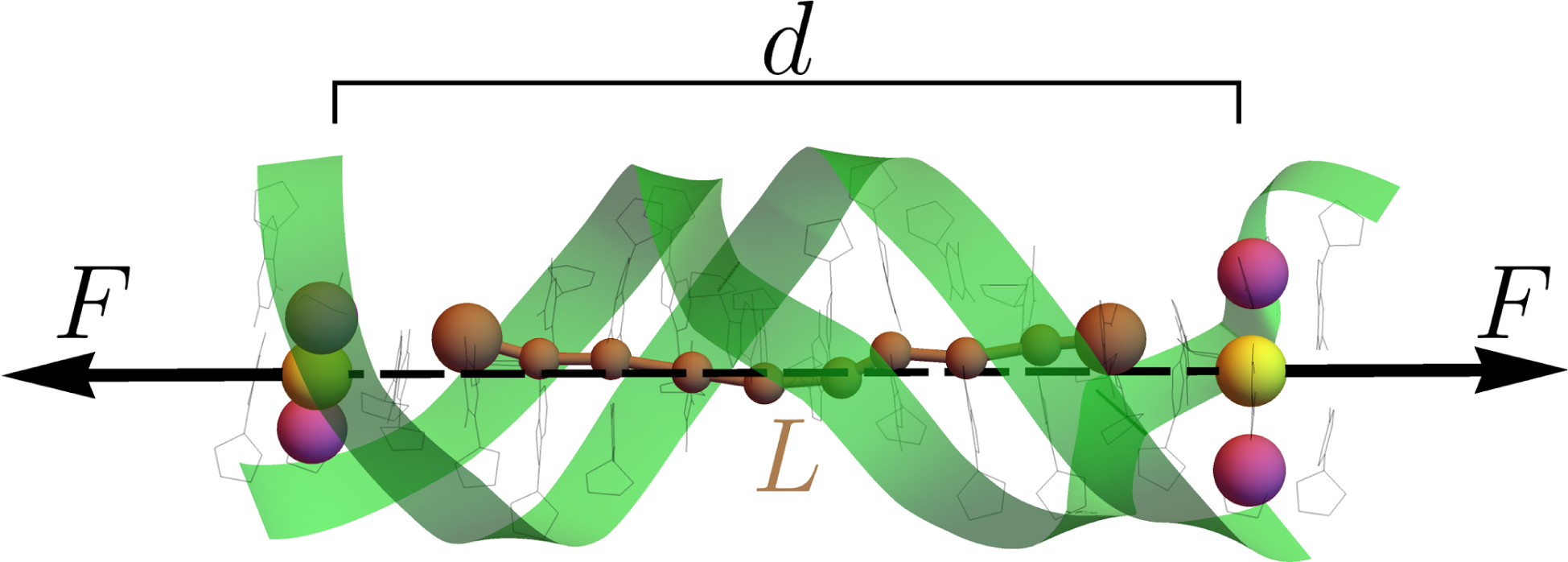
Representative snapshot illustrating the pulling protocol. Two opposite forces of magnitude *F* are applied on the centers of mass (yellow beads) of the C1’ atoms (magenta beads) of the second and second-to-last base pairs, separated by a distance *d*. The forces are aligned along the axis which connects the two centers. The overall elongation *L* is defined as the sum of helical rises along the ten central base pairs, corresponding to the contour length of the brown zig-zag line depicted in the figure. The backbone of the duplex is represented as green ribbons.

### 2.2 Analysis

The software CPPTRAJ^55^ was used to extract the structural parameters according to the 3DNA definition^56^. From them, we identified the overall extension *L* as the sum of the helical rises and the global torsion *θ* as the sum of the helical twists. In all cases, the handles were discarded from the analysis, so that only the ten central base pairs were considered (Fig. 1). As shown in Fig. S1 in the Supplementary Material for some representative cases, the large simulation time ensured a nice convergence of these variables.

In order to characterize the elasticity of dsDNA, we model it as an elastic rod which can be stretched and twisted. Thermally-induced bending is neglected as its effects can be safely ignored thanks to the combination of short sequences and applied forces employed here^38^. The energy of the system thus reads

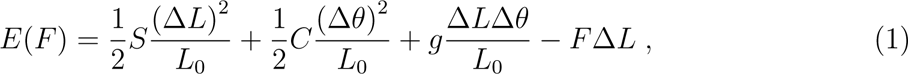

where Δ*L* = *L* − *L*_0_ and Δ*θ* = *θ* − *θ*_0_, with *L*_0_ and *θ*_0_ referring to the equilibrium values of *L* and *θ* in the absence of mechanical stress, while *F* is the value of the pulling force.

#### 2.2.1 Free-Energy Perturbation

Since we are interested in the elastic properties of the central fragment, in Eq. (1) we conjugate the force *F* with the change of the extension Δ*L*, i.e. of the contour length of the zig-zag line represented in Fig. 1. However, the simulation protocol is actually applying a couple of constant forces along the line joining the centers of the second and second-to-last base pairs (yellow beads in Fig. 1), so that in the simulations the variable conjugated to *F* is their euclidean distance *d*. Hence, the energy *Ē* regulating the conformational space being explored in the simulations is given by *Ē*(*F*) = *E*(*F*) − *F* (Δ*d* − Δ*L*). In order to properly sample the ensemble corresponding to Eq. (1), we thus employ the Free-Energy Perturbation (FEP) technique^57^. For any observable *O*, its average value ⟨*O*⟩*_F_* in the ensemble corresponding to the energy *E*(*F*) can be written as

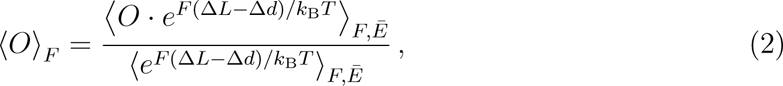

where *k*_B_*T* is the thermal energy and ⟨*. . .* ⟩_*F,Ē*_, denotes averaging in the ensemble correspond-ing to the energy *S̃*(*F*). Note that the practical application of the FEP technique relies on the assumption of a significant overlap between the conformational spaces corresponding to *E* and *Ē*. In the present case, this is indirectly supported by the observation that the widths of the distributions of *L* and *θ* are significantly larger than the shift introduced by the application of different pulling forces (see Fig. S1 in the Supplementary Material for a representative example), thus suggesting that a similar overlap occurs in the ensembles of *E*(*F*) and *Ē*(*F*) at a given force.

In practice, Eq. (2) implies computing the averages from the simulations by weighting each snapshot according to the Boltzmann weight exp [*F* (Δ*L* − Δ*d*)*/k*_B_*T*]. For each observ-able and for the corresponding fluctuations, the error was estimated by block analysis^57^. For each block size, the error associated to the sample was computed by bootstrap with 1000 repetitions, where each element was extracted with probability proportional to the corresponding weight. The weight of a block was computed by summing the FEP Boltzmann weights of the snapshots included therein. An example of the error estimation according to block size is reported in Fig. S2 in the Supplementary Information. The final value of the error was obtained by averaging over the last 200 block sizes.

#### 2.2.2 Effective stretch modulus and crookedness stiffness

The average change in extension ⟨Δ*L*⟩*_F_* can be computed from Eq. (1), leading to^38^ ⟨Δ*L*⟩*_F_* = *L*_0_*F/S̃*, where *S̃* = *S* − *g*^2^*/C* is the effective stretch modulus. Assuming *S̃* to be independent of the pulling force, its value can thus be determined from the slope of ⟨Δ*L*⟩*_F_* as a function of *F* .

The crookedness *β* is a structural parameter quantifying the displacement of the base-pair centers from the helical axis, and is defined as cos *β* = *L/* Σ*_i_ u_i_*, where *u_i_* is the center-to-center distance between base pairs *i* and *i* + 1, and the sum runs over the fragment being analysed^39^. The crookedness stiffness *k_β_* describes the response of *β* to the pulling force and is defined in analogy to the effective stretch modulus as ⟨cos *β*⟩*_F_* = *c*_0_(1 + *F/k_β_*), where *c*_0_ is the extrapolation of ⟨cos *β*⟩*_F_* at zero force.

The values of *S̃* and *k_β_* were computed by performing a fit of ⟨Δ*L*⟩*_F_* and ⟨cos *β*⟩*_F_* vs *F* according to the equations above. The associated errors were obtained as, where 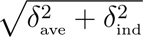, *δ*_ave_ is the error arising from the standard least-square minimization applied to the average values and *δ*_ind_ is the error originated from the indeterminacies of the various points being fitted, estimated according to the protocol described in Ref.^58^. This approach was also used in all the other fits performed in this work.

The values of *k_β_* obtained for the various systems show an approximately exponential behavior when plotted as a function of ⟨*β*⟩_1_. The function *k_β_*(⟨*β*⟩_1_) was thus fitted according to the formula *Ae*^−^*^D^*^·⟨^*^β^*^⟩1^ + *B*, where *A, B* and *D* are the adjustable parameters^39^. According to a model proposed in the literature^39^, the overall stretching response regulated by *S̃* can be ascribed to the combined effect of base-pair centers alignment (decreased crookedness) and stretching of the center-to-center distance *u_i_*:

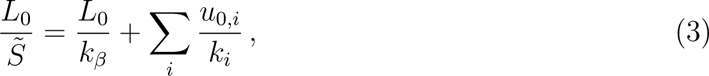

where *k_i_* and *u*_0_*_,i_* are the elastic constant and zero-force extrapolation determining the re-sponse of ⟨*u_i_*⟩*_F_* to the pulling force: ⟨*u_i_*⟩*_F_* = *u*_0_*_,i_*(1 + *F/k_i_*). By following the same procedure as in Ref.^39^, we determined the values of *k_i_* and *u*_0_*_,i_* for the ten different kinds of steps (Tables S1 and S2 in the Supplementary Material). The predicted value of *S̃* was finally computed by means of Eq. (3), where *k_β_* was obtained from ⟨*β*⟩_1_ via the exponential fit, and the associated indeterminacy estimated by error propagation.

Finally, based on Eq. (3), the contribution to the effective stretching stemming from the crookedness can be estimated as (*L*_0_*/k_β_*)*/*(*L*_0_/*S̃*) = *S̃/k_β_*.

#### 2.2.3 Computation of force-dependent elastic parameters

Following the theory presented in Ref.^41^, the elastic parameters in Eq. (1) can be computed at each value of *F* by analyzing the fluctuations involving *L* and *θ*, thus unveiling the presence of force dependence of the elastic response. For completeness, we report here the formulas employed in this work and derived in Ref.^41^:

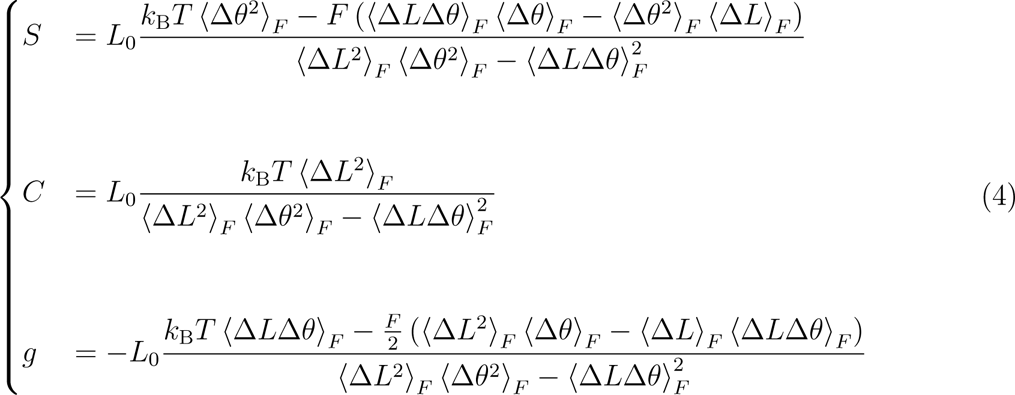

To succinctly characterize the evolution of the elastic parameters with the external force, we perform the linear fits *S*(*F*) = *S*_0_+*F* ·*dS/dF*, *C*(*F*)+*C*_0_+*F* ·*dC/dF* and *g*(*F*) = *g*_0_+*F* ·*dg/dF*, where the error on the adjustable parameters is computed as in Section 2.2.2.

## 3 Results and discussion

We performed simulations employing either the bsc1 or the OL15 modifications for a set of seven sequences, as reported in the Methods. The sequence RNG contains all possible combinations of base-pair steps, and has been studied in the past to mimic the behavior of long, random sequences employed in single-molecule experiments^38,45^. The other six sequences are obtained as pentamers of the steps AA, AC, AG, AT, CG or GG, which encompass the ten distinct base-pair steps^39^. We refer to these sequences as poly-XY, where XY is any of the six steps employed to build the pentamers.

For each sequence, after standard minimization and equilibration phases, we performed pulling simulations where two opposite, constant forces are applied to the ends of the molecule (see Methods). We run simulations at different magnitudes of such forces, namely 1, 5, 10, 15 and 20 pN. For the whole set, two external handles were added to the fragment of interest, so as to minimize end effects. In all the subsequent analysis, only the central 10 base pairs were considered. Note that this same set of sequences has been studied in the past with the bsc0 modification and with the same protocol^38,39^, enabling a direct comparison between the three different force fields in the same context.

### 3.1 The effective stretch modulus depends on the force field

The first elastic parameter investigated is the effective stretch modulus *S̃*, which can be obtained from the slope of the average change in elongation ⟨Δ*L*⟩*_F_* as a function of the applied force *F* (Section 2.2.2). As a representative case, in Fig. 2a we report the stress-vs-strain curve for the sequence RNG and the three different force fields. The slope of the curve is equal to 1/*S̃*, indicating a softer response in the case of bsc0 with respect to the other modifications. In Fig. 2b and in Table 1, we report the results obtained throughout the whole set. In the figure, the shaded grey region corresponds to the experimental values obtained in the literature in pulling experiments of large molecules, where a similar range of forces has been applied.

**Figure 2:**
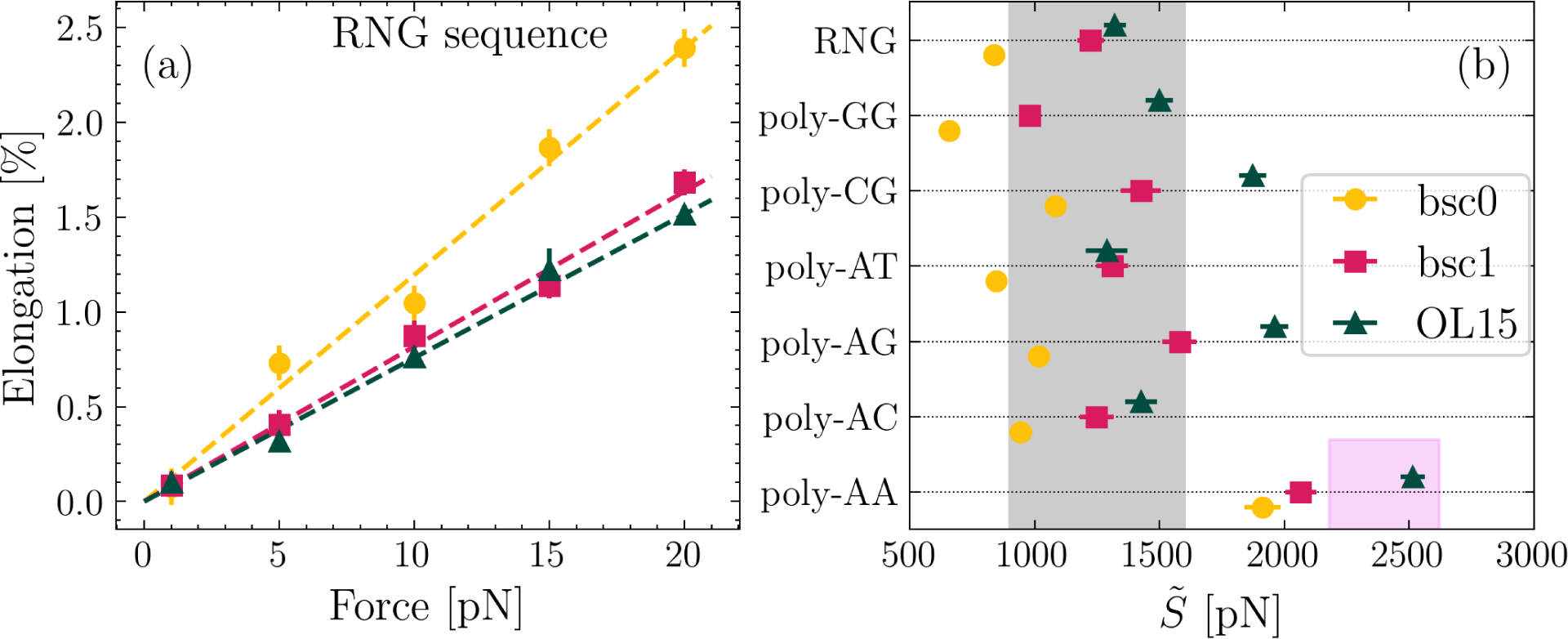
(a) Representative elongation vs force curve for the RNG sequence. The slopes of the linear fits (dashed lines) correspond to *S̃*^−1^. (b) Values of *S̃* simulated for each sequence and force field. The shaded regions indicate the experimental range for random sequences^13–16^ (gray) and phased A-tracts^9^ (pink).

**Table 1:**
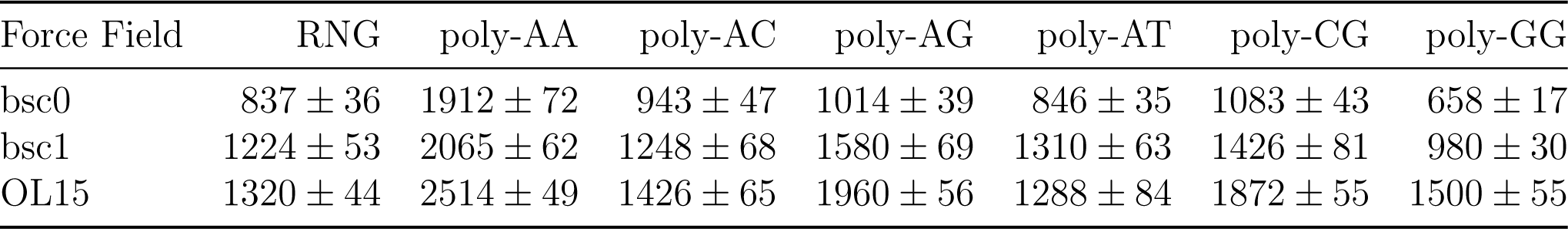
Values of *S̃* in pN for the various sequences and force fields.

From Fig. 2b and Table 1, a clear pattern emerges, according to which the values of *S̃* are systematically ranked in the order bsc0 < bsc1 < OL15, thus outlining a clear difference in the stiffness predicted by the various force fields. At a general level, the values obtained for bsc0 and bsc1 mostly lie within the experimentally-known range, while for OL15 they are usually found to be larger. While this suggests that dsDNA within OL15 might be too stiff, one should take this indication with some caution, as the sequences poly-XY are not directly comparable with long, random sequences usually employed in single-molecule experiments. In this regard, the most valuable system is provided by the random-like sequence RNG, for which the three force fields predict values of *S̃* within the experimental range. Interestingly, in all cases the sequence poly-AA is found to be significantly stiffer than the other molecules. The most direct experimental comparison for this system is provided by the phased A-tracts reported in a recent study^9^, where pulling by optical tweezers estimated *S̃* = (2400±220) pN. This is in line with the three predictions, with the best quantitative agreement being found for OL15.

### 3.2 Stretching is determined by crookedness

The crookedness *β* is a structural parameter characterizing the displacement of the base-pair centers from the helical axis^39^. By definition, larger values of *β* indicate a more crooked structure, with *β* = 0 corresponding to perfectly aligned centers (see Methods for the formal definition). It was proposed that the elongation response of dsDNA to a pulling force can be described as the combined effect of force-induced alignment of the centers and stretching of the center-to-center distance between consecutive base pairs^39^. Intriguingly, this led to a model which, in the case of bsc0, quantitatively predicted the value of *S̃* from knowledge of *β* in the absence of mechanical stress, combined with tabulated values of stiffness of the ten possible base-pair steps combinations. In other terms, this allows predicting the mechanical response from the sequence of the dsDNA fragment and from knowledge of its structure in the absence of applied force.

We employed the extended dataset to check whether this model is also applicable to the other force fields, or rather it is restricted to bsc0. Following Ref.^39^, we first characterized the stiffness *k_β_* associated to *β*, which accounts for the energetic cost needed to align the centers (see Methods for the formal definition). In Fig. 3a, we report the values of *k_β_* as a function of the average crookedness ⟨*β*⟩_1_ obtained at *F* = 1 pN, which we use as an estimator for the unperturbed molecule. Remarkably, the whole dataset corresponding to the three force fields can be nicely fitted by a single function of the form *Ae*^−^*^D^*^·⟨^*^β^*^⟩1^ + *B*. This phenomenological functional form was proposed in Ref. ^39^ for the bsc0 data, and is found here to be able to capture simultaneously the data for the three force fields. The values of the optimized parameters are *A* = (0.29 ± 0.07) · 10^6^ pN, *D* = 12.8 ± 0.7 and *B* = (736 ± 37) pN. This emerging pattern suggests that the difference in rigidity of the various modifications can be ultimately ascribed to the improved structural features introduced in bsc1 and OL15, at least at the level of the crookedness stiffness.

**Figure 3:**
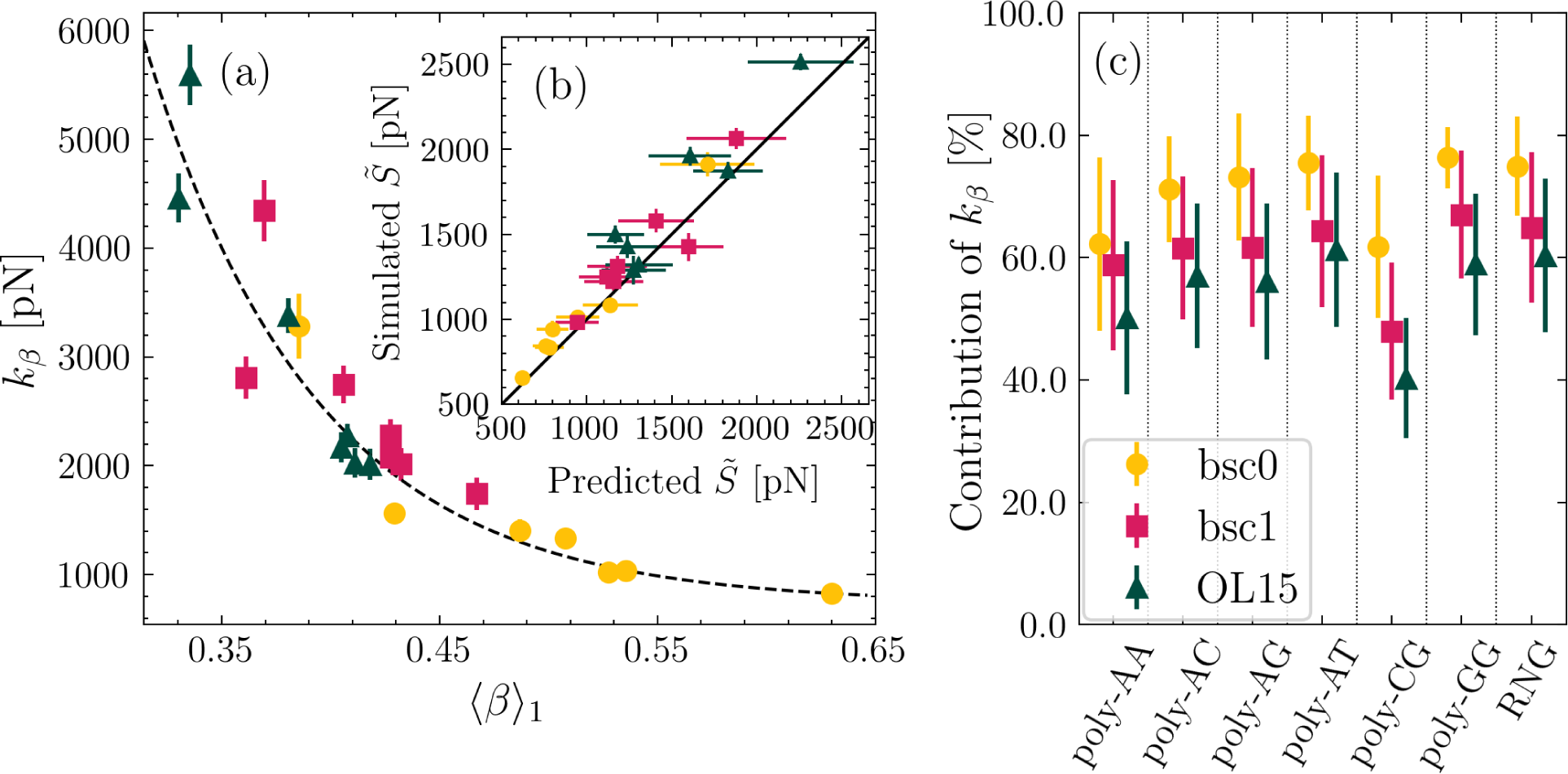
(a) Elastic constant associated with crookedness, *k_β_*, as a function of the average crookedness at 1 pN, ⟨*β*⟩_1_. Dashed line is a fit to the function *k_β_* = *Ae*^−^*^D^*^·⟨^*^β^*^⟩1^ + *B*. (b) Comparison between predicted values of *S̃* and results obtained from simulations. (c) Con-tribution of *k_β_* to the overall stretching response.

As the next step, we employed the simulation results for poly-XY to determine the elastic constants associated to the ten distinct base-pair step combinations ^39^ (reported in Table S1 in the Supplementary Material). In contrast with the case of *k_β_*, inspection of the step stiffness constants did not reveal a clear pattern in their dependence on the chosen force field. This further supports the idea that it is the change in structure (i.e. the unperturbed crookedness) which ultimately originates the ranking bsc0 < bsc1 < OL15 observed for *S̃* in Fig. 2b and Table 1.

Building on this point, we consider the model from Ref.^39^, which predicts the value of *S̃* by considering an effective series of springs involving the crookedness rigidity and the center-to-center distances (see Methods). The value of *k_β_* is computed from the crookedness of the unperturbed system (here estimated as ⟨*β*⟩_1_) via the empirical exponential function. As for the steps, given the absence of a clear force-field dependence, and with the aim of building a theoretical framework including the three modifications at the same time, we consider for the elastic parameters the averages of the values obtained for bsc0, bsc1 and ol15 (see Section S2 in the Supplementary Material). In Fig. 3b, we compare *S̃* as predicted by the model and the values retrieved directly from the simulations. The close agreement between prediction and numerical results further endorses the universality of the model across the spectrum of the different force fields.

Based on the model, we further investigated the contribution of *k_β_* to *S̃*^39^, which was estimated as *S̃/k_β_* (see Methods) and is reported in Fig. 3c. In most cases, the crookedness is the major responsible for the response to the pulling force^39^, although from a quantitative perspective its contribution is lower for bsc1 and OL15. This is in line with the absence of a pattern in the step stiffness constants which, once combined with the stiffening of *k_β_* induced by the enhanced spontaneous alignment of the centers, results into an overall smaller contribution of the crookedness stiffness.

### 3.3 Stretch modulus increases with force

The effective stretch modulus *S̃* quantifies the net elongation response to the pulling force. Due to a coupling between the extension *L* and the torsion *θ* of the duplex, the net elongation change ⟨Δ*L*⟩*_F_* induced by *F* results from the interplay between these two degrees of freedom. Quantitatively, one finds that *S̃* = *S* − *g*^2^*/C*, where *S* is the stretch modulus, *C* is the twist modulus and *g* is the twist-stretch coupling (see Methods for further details).

Nevertheless, the description of dsDNA via only two variables is necessarily an effective one, so that the elastic parameters *S*, *C* and *g* are expected to change according to the applied mechanical stress^41^. We have recently introduced a theoretical framework which allows to determine the force-dependent values of the elastic parameters by studying their fluctuations in the constant-force ensemble and employed it to analyze the force-dependent behavior of dsDNA and dsRNA^41^. For dsDNA, in our previous work we employed the bsc0 data. Here, we study the extended dataset to check the robustness of our conclusions with respect to the force field considered. For completeness, we report in the Methods the formulas derived in Ref.^41^ which are employed in the present study (Eq. (4)).

In Fig. 4a, we report the values of *S* obtained by fluctuations at the smallest force *F* = 1 pN. Given the low value of *F*, they provide a good approximation of the unperturbed case. The data show a strong resemblance with the values of *S̃* reported in Fig. 2b. This is expected, since the typical values of *g* and *C* (see below) provide a small correction when employing the formula *S̃* = *S* − *g*^2^*/C*, thus leaving unchanged the pattern observed above, i.e. the systematic ranking bsc0 < bsc1 < OL15. For completeness, we also report the values of *S* published in the literature and computed by means of fluctuations in the absence of a pulling force^40,48,51^. Particularly, the data labeled as “Dohnalova2022” consider the same sequence for bsc1 and OL15^40^, confirming for a specific case our general observation on the higher rigidity of OL15.

**Figure 4:**
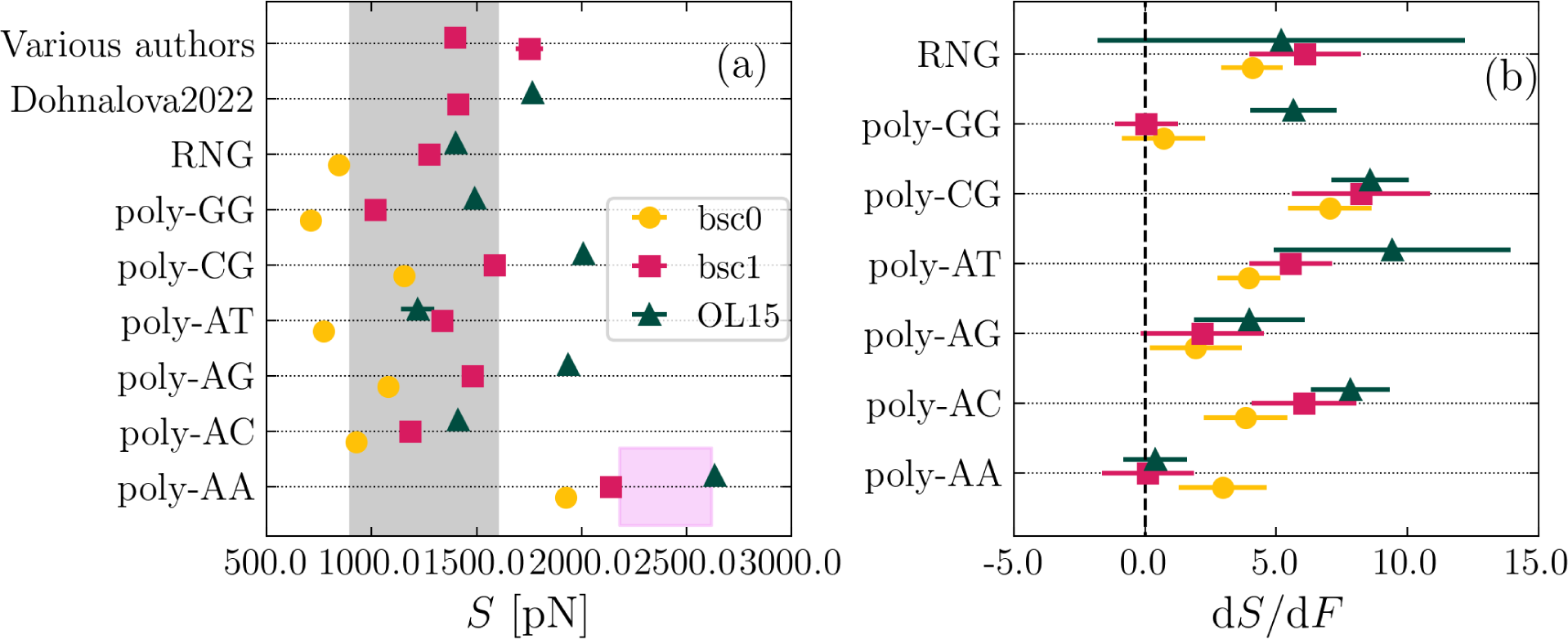
(a) Values of *S* at 1 pN computed by analysis of fluctuations. The shaded region corresponds to the experimental range. Due to the lack of direct experimental determination of *S*, the shaded region corresponds to the values of *S̃* measured in the literature, as in Fig. 2b. (b) Slope of the *S*(*F*) curve obtained for the various sequences and force fields.

To characterize the force-induced change in the stretch modulus, we performed a linear fit of *S*(*F*), for which the slope *dS/dF* is reported in Fig. 4b. For all sequences, *dS/dF* ≥ 0, indicating that dsDNA stiffens upon pulling, in agreement with our previous report on bsc0^41^. Microscopically, this stiffening can be ascribed to the progressive alignment of the aromatic rings of consecutive base pairs due to the action of the force, which increases the strength of stacking interactions^41^. This picture is supported quantitatively by the significant correlation (*r* ≃ 0.70) between the change in stretch modulus *dS/dF* and the change in slide *dλ/dF*, as can be observed from Fig. S3 in the Supporting Material. All in all, the present analysis fully extends to the bsc1 and OL15 force fields our previous findings on the force-induced stiffening of dsDNA^41^.

### 3.4 Twist modulus ranking depends on sequence

Our results for the twist modulus *C* are reported in Fig. 5. In Fig. 5a, we show the values of *C* obtained at *F* = 1 pN, together with data collected from the literature^40,48,50,51^. The agreement with the experimental values (shaded area) is generally good for all the force fields, particularly when considering random-like sequences (first four rows in Fig. 5a). The ranking bsc0 < bsc1 < OL15 holds for several sequences (poly-AA, poly-AG, poly-CG, RNG), but is not as general as we found for *S*. For instance, for poly-AT OL15 is found to be as soft as bsc0, while bsc0 shows the highest twist stiffness in the case of poly-AC. The most peculiar case is provided by poly-GG, for which bsc0 has a much larger value of *C* than both the other modifications, despite showing enhanced flexibility in the stretching mode (Fig. 4a). This case reflects previous observations on the dependence of rigidity on the elastic mode under consideration^59^. The lack of a general pattern in *C* is supported by other data from the literature, which show reversed rankings for bsc1 and OL15 (OL15 < bsc1 in Ref.^40^, bsc1 ≲ OL15 for the 32mer in Ref. ^50^).

**Figure 5:**
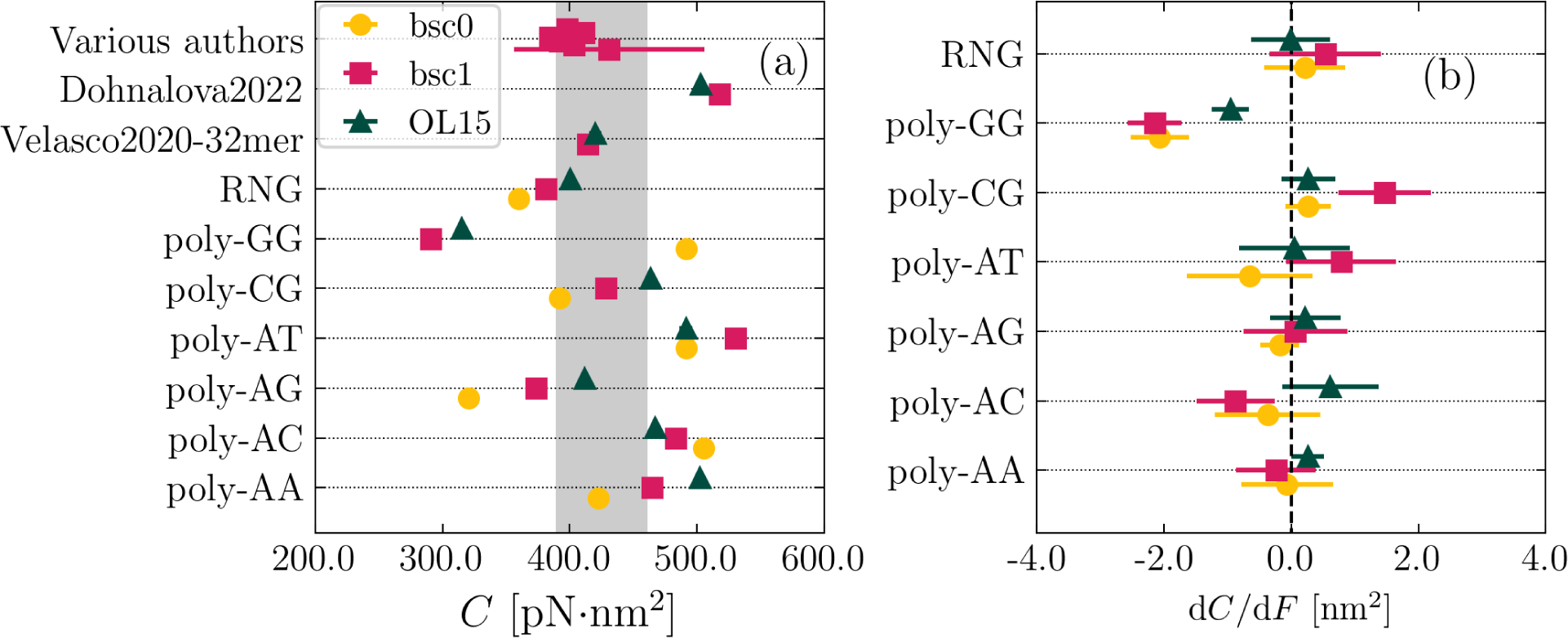
(a) Values of *C* at 1 pN computed by analysis of fluctuations. The shaded region corresponds to the experimental range. (b) Slope of the *C*(*F*) curve obtained for the various sequences and force fields.

In Fig. 5b, we report the slope *dC/dF* to characterize the force dependence of *C*. We observe virtually no change in the twist modulus, with the exception of the sequence poly-GG, which shows a clear negative slope for all cases. It is worth mentioning that this duplex is the most crooked molecule, for which the force-induced straightening of the spontaneous curvature is likely to induce a softer twist response^41^.

### 3.5 The three modifications qualitatively capture the twist-stretch coupling

In Fig. 6 we report the results obtained for the twist-stretch coupling *g*. In all cases, we find that *g <* 0 (Fig. 6a), in agreement with experimental results^19^. Quantitatively, there is a tendency of all the force fields to overestimate the magnitude of *g* with respect to experiments (shaded region in Fig. 6a), as observed in previous work^38^. While this feature might be a potential target to address in future refinements of the force fields, such overestimation must be taken with caution, due to the likely dependence of *g* on the length of the fragment under consideration^58^. Particularly, based on a nucleotide-level coarse-grained model, we have previously reported significant changes of *g* in the 20-40 base-pairs range^58^, so that such an effect is expected to be further enhanced for the large molecules employed in experiments. In terms of ranking, in analogy to *C* no clear patterns are present, although one may notice a generic shift of OL15 values towards larger magnitudes of *g*.

**Figure 6:**
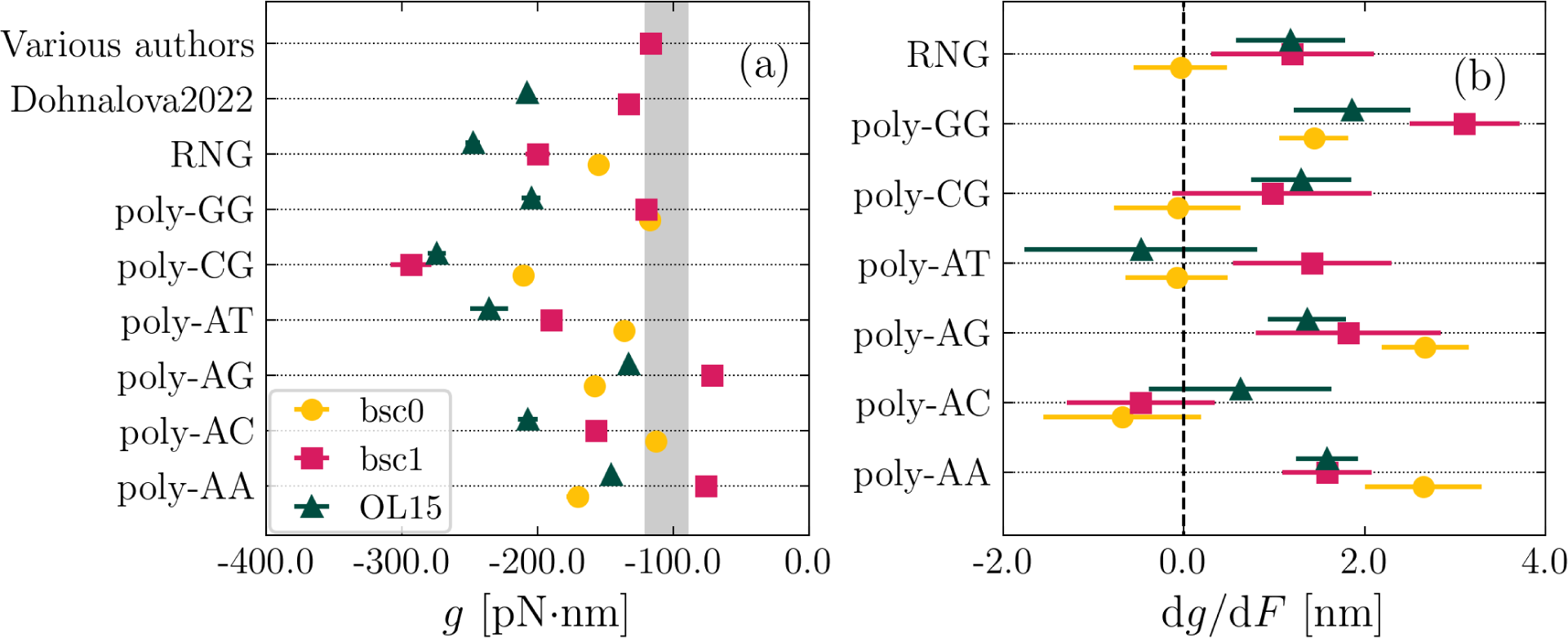
(a) Values of *g* at 1 pN computed by analysis of fluctuations. The shaded region corresponds to the experimental range. (b) Slope of the *g*(*F*) curve obtained for the various sequences and force fields.

In Fig. 6b, we report the slope *dg/dF* characterizing the change of twist-stretch coupling with force. For all force fields and all sequences, one has *dg/dF* ≥ 0 within error, indicating a weakening of *g* with the pulling force. Again, this is in line with previous reports on bsc0^38^ and with the experimental literature^19,21^. As a future direction, it will be an interesting challenge to check the performance of the various force fields in capturing the experimentally-observed sign reversal of *g* at forces around 40 pN.

## Conclusions

In this work, we have performed a systematic comparison of the molecular mechanics of dsDNA as predicted by the most advanced force fields for dsDNA based on the parm99 model, bsc1 and OL15, as well as their common predecessor bsc0. We have found that the global response of the duplex to a pulling force is captured with good precision by all the models, although some clear differences are evidenced.

The three modifications show similar behavior in the twist modulus and in the twist-stretch coupling, in terms of both their average values and their evolution with the magnitude of the external force being applied. As for the stretching, a clear pattern is observed, according to which the force fields are ranked from the softest to the stiffest as bsc0 < bsc1 < OL15. This feature is perhaps the most prominent difference observed to date between the bsc1 and OL15 modifications, for which previous studies have reported very similar performance in reproducing the structural features of dsDNA^49^. Intriguingly, we could ascribe this difference in mechanics to the values of the crookedness predicted by the force fields in the absence of mechanical stress, extending the reach of previous observations^39^ and unveiling a common mechanism originating the different stretching response of the three modifications.

Overall, the differences observed for the stretching modulus suggest that bsc1 might be preferred over OL15 for studies focused on the mechanics of dsDNA. However, due to the peculiar features of the poly-XY sequences from which this difference is argued, this observation has to be taken with caution. In order to conclusively determine the force field with the best performance, it would be ideal to couple the present results with single-molecule experiments on large poly-XY sequences, so as to provide the ground for a thorough and decisive comparison.

## Acknowledgement

The project that gave rise to these results received the support of a fellowship from ‘la Caixa’ Foundation (ID 100010434) and from the European Union’s Horizon research and innovation programme under the Marie Skłodowska-Curie grant agreement No. 847648. The fellowship code is LCF/BQ/PI20/11760019. We acknowledge support from the Ministerio de Ciencia e Innovación (MCIN) through the project PID2020-115864RB-I00 and the ‘María de Maeztu’ Programme for Units of Excellence in R&D (grant No. CEX2018-000805-M). The authors thankfully acknowledge the computer resources at MinoTauro and Finisterrae3 and the technical support provided by Barcelona Supercomputing Center (RES-FI-2021-2-0042, RES-FI-2022-1-0016).

