## Supplemental Information for "Systematic comparison of Amber force fields for the mechanical properties of double-stranded DNA"

### 1 Convergence of simulations

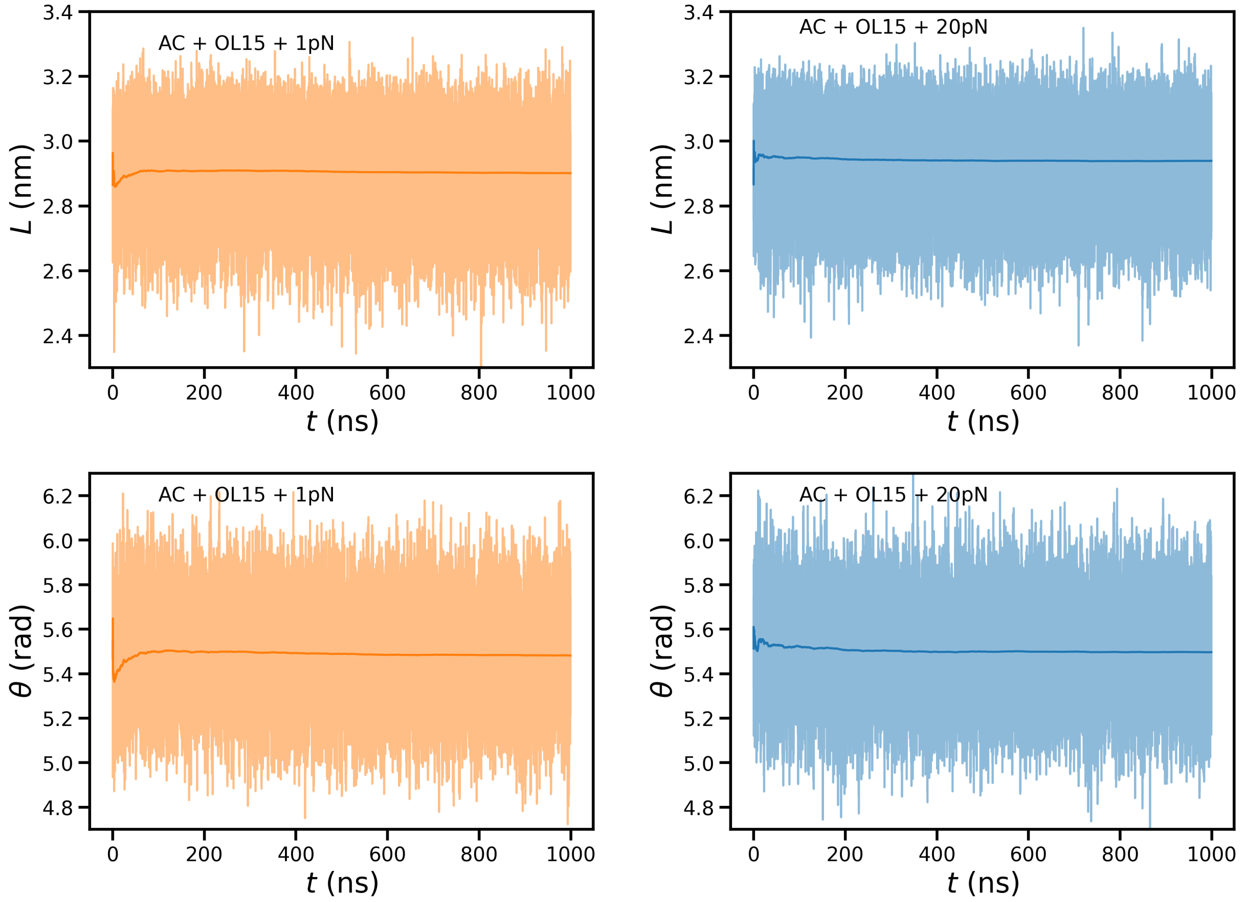


Figure S1: Representative examples of convergence for the extension $L$ (top) and the torsion $\theta$ (bottom). Transparent lines show the trajectories $L(t)$ and $\theta(t)$, with $t$ being the simulation time. Full lines report the running average up to time $t$ along the simulation time. Left and right column correspond to pulling forces equal to $F = 1$ pN and $F = 20$ pN, respectively. From their comparison, one can appreciate the different magnitude between the force-induced shift of the average value and the thermally-induced fluctuations at each force.

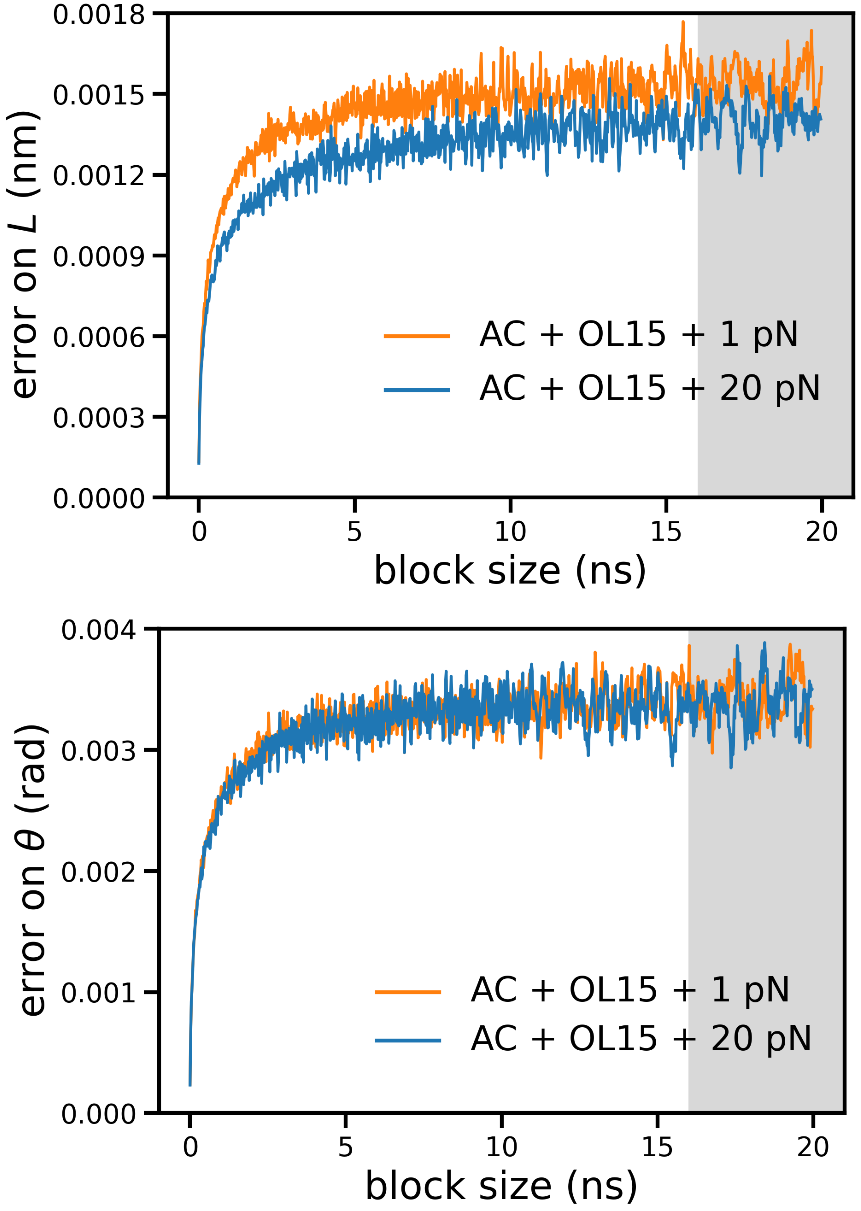


Figure S2: Representative examples of block analysis for error estimation on $L$(top) and $\theta$ (bottom). The final error was obtained by averaging the values of the last 200 block sizes, highlighted by the shaded region.

#### 2 Step-dependent stretching stiffness

Table S1: Elastic constant $k$ and unperturbed length $u_0$ for the 10 different kinds of steps and the various force fields. For each case, the values were determined by collecting the force-dependent average extension $\langle u \rangle_F$ of all the steps of the kind under inspection from the sequences poly-XY. Then, the parameters were obtained by fitting the results via the formula $\langle u \rangle_F = u_0(1 + F/k_i)$. In the case of $u_0$, errors are not listed as they are smaller than the last reported figure. For some steps, the constant $k$ is characterized by large (possibly negative) values and high error, highlighting in practice no force dependence for $u$ (e.g. see step GA).

Force-field dependence

| | bsc0 | | bsc1 | | OL15 | |
|---|---|---|---|---|---|---|
| Step | $k$ (pN) | $u_0$ (nm) | $k$ (pN) | $u_0$ (nm) | $k$ (pN) | $u_0$ (nm) |
| AA | $4865 \pm 365$ | 0.352 | $4066 \pm 393$ | 0.352 | $4766 \pm 227$ | 0.346 |
| AC | $1856 \pm 333$ | 0.353 | $3561 \pm 450$ | 0.351 | $4450 \pm 567$ | 0.341 |
| AG | $2226 \pm 189$ | 0.367 | $2419 \pm 307$ | 0.357 | $3401 \pm 350$ | 0.356 |
| AT | $3187 \pm 261$ | 0.347 | $4061 \pm 581$ | 0.340 | $5599 \pm 1254$ | 0.336 |
| CA | $8137 \pm 3657$ | 0.355 | $2434 \pm 336$ | 0.365 | $4595 \pm 697$ | 0.364 |
| CG | $2721 \pm 408$ | 0.339 | $1903 \pm 144$ | 0.346 | $2678 \pm 132$ | 0.355 |
| GA | $-31176 \pm 28986$ | 0.354 | $19164 \pm 15880$ | 0.358 | $13811 \pm 4640$ | 0.347 |
| GC | $7610 \pm 1532$ | 0.359 | $25961 \pm 23327$ | 0.356 | $4785 \pm 572$ | 0.343 |
| GG | $3355 \pm 207$ | 0.379 | $2319 \pm 226$ | 0.369 | $5811 \pm 821$ | 0.359 |
| TA | $10875 \pm 3628$ | 0.363 | $3512 \pm 575$ | 0.374 | $2702 \pm 442$ | 0.374 |

Table S2: Elastic constant $k$ and unperturbed length $u_0$ for the 10 different kinds of steps averaged over the three force fields. For each case, the values were determined from Table S1 by considering the weighted average $w_{k,\mathrm{bsc0}}k_{\mathrm{bsc0}} + w_{k,\mathrm{bsc1}}k_{\mathrm{bsc1}} + w_{k,\mathrm{OL15}}k_{\mathrm{OL15}}$ and $w_{u,\mathrm{bsc0}}u_{0,\mathrm{bsc0}} + w_{u,\mathrm{bsc1}}u_{0,\mathrm{bsc1}} + w_{u,\mathrm{OL15}}u_{0,\mathrm{OL15}}$. In the previous formula, each weight is proportional to the inverse of the squared error from Table S1, and the normalizations $w_{k,\mathrm{bsc0}} + w_{k,\mathrm{bsc1}} + w_{k,\mathrm{OL15}} = 1$ and $w_{u,\mathrm{bsc0}} + w_{u,\mathrm{bsc1}} + w_{u,\mathrm{OL15}} = 1$ hold. Errors were propagated from Table S1 as $\sqrt{\delta_1^2 + \delta_2^2}$ where, in the case of $k$, $\delta_1 = \sqrt{(w_{k,\mathrm{bsc0}}\delta k_{\mathrm{bsc0}})^2 + (w_{k,\mathrm{bsc1}}\delta k_{\mathrm{bsc1}})^2 + (w_{k,\mathrm{OL15}}\delta k_{\mathrm{OL15}})^2}$ and $\delta_2$ is the weighted standard deviation of the values of $k$. A similar formula applies for $u_0$.

| Average values | | |
|---|---|---|
| Step | $k$ (pN) | $u_0$ (nm) |
| AA | $4653 \pm 337$ | $0.348 \pm 0.003$ |
| AC | $2821 \pm 1092$ | $0.346 \pm 0.005$ |
| AG | $2475 \pm 457$ | $0.360 \pm 0.005$ |
| AT | $3413 \pm 575$ | $0.343 \pm 0.004$ |
| CA | $2877 \pm 995$ | $0.362 \pm 0.004$ |
| CG | $2345 \pm 398$ | $0.352 \pm 0.005$ |
| GA | $13185 \pm 8239$ | $0.351 \pm 0.004$ |
| GC | $5142 \pm 1172$ | $0.351 \pm 0.007$ |
| GG | $2982 \pm 746$ | $0.372 \pm 0.009$ |
| TA | $3076 \pm 917$ | $0.367 \pm 0.005$ |

#### 3 Relation between changes in stretch modulus and slide

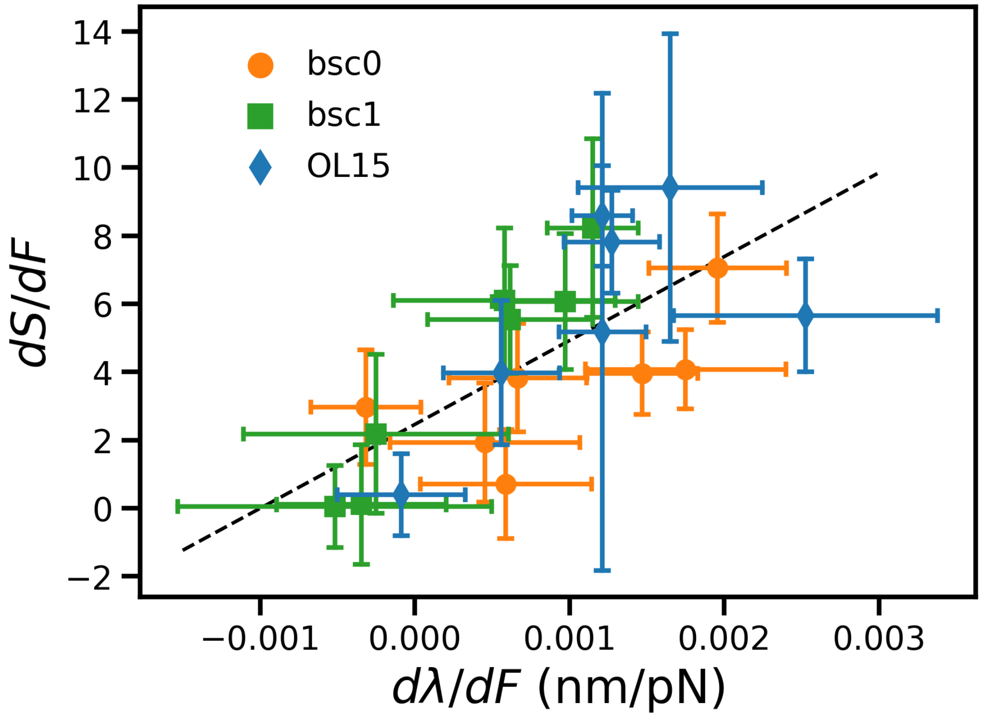


Figure S3: Variation $dS/dF$ of stretch modulus and of slide $d\lambda/dF$ upon changing the magnitude of the pulling force $F$. The dashed line represents a linear fit of the whole dataset.
